## Supplemental Figures and Tables for "Exploring the prokaryote-eukaryote interplay in microbial mats from an Andean athalassohaline wetland"

Table S1. Location and date of sampling and physicochemical parameters measured in the water layer of the sampling poin

| Site | Coordinates | Date | Temp (°C) | pH | Cond (µS/cm) | TDS (mg/l) | Salinity PSU | $O_2$ (%) |
|---|---|---|---|---|---|---|---|---|
| H6-1 | 20° 19' 53.1'' N<br>68° 50' 18.6'' W | 07.12.2019 | 21.9 | 8.82 | 1464 | 1464 | 0.7 | 107.3 |
| H6-2 | 20°20'1.5'' N<br>68°50'17.8'' W | 07.12.2019 | 24.7 | 8.57 | 2350 | 2330 | 1.2 | 212 |
| H6-3 | 20°20'3.9'' N<br>68°50'18.4'' W | 07.12.2019 | 18.5 | 7.05 | 2060 | 2270 | 1.1 | 42.3 |
| H6-4 | 20°20'7.1'' N<br>68°50'19.2'' W | 07.12.2019 | 18 | 9.23 | 1039 | 829 | 0.5 | 150 |
| H4-1 | 20°17'29.1'' N<br>68°53'18.3'' W | 08.12.2019 | 5.3 | 8.94 | 31400 | 31400 | 18.9 | 2.6 |
| H4-2 | 20°17'28.9'' N<br>68°53'18.9'' W | 08.12.2019 | 6.5 | 8.79 | 1958 | 1984 | 0.9 | 0 |
| H4-3 | 20°17'29.6'' N<br>68°53'23.5'' W | 08.12.2019 | 10.9 | 8.37 | 2020 | 2000 | 1 | 2.2 |
| H4-4 | 20°17'29.6'' N<br>68°53'23.5'' W | 08.12.2019 | 12.9 | 8.6 | 2420 | 2360 | 1.2 | 36.5 |
| H3-1 | 20°17'0.4'' N<br>68°53'19.5'' W | 08.12.2019 | 17.1 | 8.08 | 570 | 577 | 0.2 | 113.4 |
| H3-3 | 20°17'0.4'' N<br>68°53'19.5'' W | 08.12.2019 | 22.1 | 8.92 | 1242 | 1120 | 0.6 | 75.5 |
| H0-1 | 20°15'51.4'' N<br>68°52'27.9'' W | 08.12.2019 | 20.9 | 8.92 | 588 | 588 | 0.2 | 192.2 |

Table S2. Samples analyzed, sequence statistics and diversity indexes. ASV, amplicon sequence variant. Sediment always refers to sediment layers below the corresponding microbial mats

| Sample name | Site | Nature | 16S rRNA amplicon sequences | | 18S rRNA amplicon sequences | | Number of ASVs | | | Diversity indexes (prokaryotes) | | | | | Diversity indexes (eukaryotes) | | | | |
|---|---|---|---|---|---|---|---|---|---|---|---|---|---|---|---|---|---|---|---|
| | | | Total | Retained high-quality | Total | Retained high- | Archaea | Bacteria | Eukaryotes | Simpson index | Chao1 | Shannon-Wiener index | Species richnes | Pielou's index | Simpson index | Chao1 | Shannon-Wiener index | Species richnes | Pielou's index |
| H0-1-M | H0 | mat | 30340 | 13692 | 42418 | 6879 | 6 | 264 | 112 | 0.92 | 270.00 | 5.45 | 270 | 0.67 | 0.90 | 112.00 | 4.84 | 112 | 0.71 |
| H0-1-S | H0 | sediment | 27666 | 12555 | 24686 | 6580 | 118 | 582 | 103 | 0.99 | 700.56 | 8.72 | 700 | 0.92 | 0.91 | 103.43 | 4.67 | 103 | 0.70 |
| H3-1-BF | H3 | biofilm | 6891 | 2703 | 11421 | 3043 | 0 | 142 | 76 | 0.97 | 142.00 | 6.06 | 142 | 0.85 | 0.96 | 112.00 | 5.16 | 76 | 0.83 |
| H3-1-S | H3 | sediment | 13105 | 6420 | 31580 | 9110 | 43 | 337 | 116 | 0.99 | 380.00 | 7.83 | 380 | 0.91 | 0.92 | 116.00 | 4.82 | 116 | 0.70 |
| H3-3-M | H3 | mat | 58888 | 38321 | 56058 | 12825 | 0 | 870 | 247 | 0.99 | 870.20 | 8.36 | 870 | 0.86 | 0.96 | 259.21 | 5.95 | 247 | 0.75 |
| H4-1-M | H4 | mat | 57718 | 39906 | 58753 | 21351 | 1 | 423 | 73 | 0.98 | 424.00 | 6.92 | 424 | 0.79 | 0.91 | 73.00 | 4.19 | 73 | 0.68 |
| H4-1-S | H4 | sediment | 41020 | 26026 | 57196 | 28986 | 12 | 698 | 101 | 0.99 | 710.00 | 8.09 | 710 | 0.85 | 0.93 | 102.00 | 4.54 | 101 | 0.68 |
| H4-2-M | H4 | mat | 35784 | 18363 | 40619 | 18046 | 1 | 565 | 95 | 0.99 | 568.77 | 8.01 | 566 | 0.88 | 0.83 | 95.00 | 3.90 | 95 | 0.59 |
| H4-2-ME | H4 | mat | 64188 | 41483 | 44406 | 14799 | 13 | 419 | 76 | 0.92 | 432.14 | 5.64 | 432 | 0.64 | 0.92 | 76.00 | 4.51 | 76 | 0.72 |
| H4-4-M | H4 | mat | 43402 | 24926 | 46074 | 17320 | 13 | 908 | 127 | 0.99 | 921.00 | 8.44 | 921 | 0.86 | 0.94 | 127.33 | 4.72 | 127 | 0.68 |
| H6-1-M | H6 | mat | 38878 | 20528 | 44488 | 11796 | 9 | 673 | 109 | 1.00 | 682.14 | 8.50 | 682 | 0.90 | 0.94 | 111.55 | 4.85 | 109 | 0.72 |
| H6-1-S_0-5cm | H6 | sediment | 24752 | 11469 | 28568 | 9548 | 33 | 523 | 61 | 0.99 | 556.93 | 8.32 | 556 | 0.91 | 0.93 | 61.20 | 4.59 | 61 | 0.77 |
| H6-1-S_5-10cm | H6 | sediment | 23915 | 11887 | 19429 | 7462 | 75 | 553 | 64 | 1.00 | 628.56 | 8.60 | 628 | 0.93 | 0.93 | 64.00 | 4.62 | 64 | 0.77 |
| H6-2-M | H6 | mat | 35624 | 24333 | 51924 | 13619 | 2 | 559 | 160 | 0.99 | 561.43 | 7.85 | 561 | 0.86 | 0.96 | 162.33 | 5.41 | 160 | 0.74 |
| H6-2-S | H6 | sediment | 15375 | 11294 | 35945 | 19849 | 18 | 708 | 159 | 1.00 | 726.00 | 9.07 | 726 | 0.95 | 0.99 | 159.00 | 7.10 | 159 | 0.97 |
| H6-3-M | H6 | mat | 55205 | 33729 | 81143 | 46990 | 16 | 511 | 108 | 0.98 | 527.19 | 6.71 | 527 | 0.74 | 0.81 | 108.75 | 3.37 | 108 | 0.50 |
| H6-3-S | H6 | sediment | 42470 | 28448 | 57018 | 25628 | 16 | 952 | 303 | 0.99 | 968.81 | 8.80 | 968 | 0.89 | 0.98 | 308.63 | 6.83 | 303 | 0.83 |
| H6-3-W-CT | H6 | water | 57470 | 43121 | 62163 | 28471 | 227 | 1494 | 144 | 0.99 | 1725.69 | 9.21 | 1721 | 0.86 | 0.90 | 145.20 | 4.80 | 144 | 0.67 |
| H6-4-M | H6 | mat | 49348 | 24199 | 69158 | 31579 | 0 | 215 | 51 | 0.88 | 215.00 | 4.33 | 215 | 0.56 | 0.87 | 51.00 | 3.39 | 51 | 0.60 |

**Table S3.** Frequency and phylogenetic affiliation of prokaryotic ASVs in Salar de Huasco samples.

**Table S4.** Frequency and phylogenetic affiliation of eukaryotic ASVs in Salar de Huasco samples.

These tables are very large and can be downloaded as an excel file.

Table S5. Parameters of co-occurrence network topologies

| | | |
|---|---|---|
| General network | Nodes | 395 |
| | Edges | 1832 |
| | Postive | 1368 |
| | Negative | 464 |
| Filtered network | Nodes | 119 |
| | Prokaryotes | 58.8% |
| | Eukaryotes | 41.2% |
| | Edges | 104 |
| | Diameter | 2.1 |
| | Mean distance | 0.68 |
| | Cluster | 19 |
| | Modularity | 0.92 |

**Table S6.** Statistics of major clusters of co-occurrent prokaryotic and eukaryotic ASVs.

| Cluster | Domain | Phylum | Class | Order | Family | Genus | Connections | Label | Presence in mat samples (%) | Presence in sediment samples (%) |
|---|---|---|---|---|---|---|---|---|---|---|
| Cluster 1 | | | | | | | | | | |
| dc318e4f93e5c7cf6166ad063ce2ce89 | Eukaryota | Ochrophyta | Bacillariophyta | Bacillariophyta_X | Raphid-pennate | NA_Raphid-pennate | 1 | 7 | 50.1 | 49.9 |
| 2bcdc0b08623c436bcbeca57eeee6612 | Eukaryota | Ochrophyta | Bacillariophyta | Bacillariophyta_X | Raphid-pennate | NA_Raphid-pennate | 1 | 7 | 50.3 | 49.7 |
| 81a1a52c4affb568a520c03348f8cdef | Prokaryote | Proteobacteria | Gammaproteobacteria | Ectothiorhodospirales | Ectothiorhodospiraceae | Ectothiorhodospira | 3 | 16 | 58.3 | 41.7 |
| 79f77063b4dc797b3bd3ed0b58a21491 | Prokaryote | Proteobacteria | Alphaproteobacteria | Rhodobacterales | Rhodobacteraceae | NA_Rhodobacteraceae | 1 | 16 | 68.2 | 31.8 |
| 766f615ebb91e5ab291122da9ab3a12f | Prokaryote | Desulfobacterota_I | Desulfovibrionia | Desulfovibrionales | Desulfomicrobiaceae | Desulfomicrobium | 2 | 13 | 74.4 | 25.6 |
| Cluster 2 | | | | | | | | | | |
| e073b562eafd267fe500fb68805c0c1b | Prokaryote | Gemmatimonadota | Gemmatimonadetes | Gemmatimonadales | Gemmatimonadaceae | Gemmatimonas | 1 | 15 | 59.9 | 40.1 |
| 38cd76508fe018b57eb5f757d3c24e2c | Prokaryote | Bacteroidota | Bacteroidia | Bacteroidales | Lentimicrobiaceae | Lentimicrobium | 1 | 9 | 24.0 | 76.0 |
| 7a5c9e45b3051c4653b7c38614f692ac | Prokaryote | Bacteroidota | Bacteroidia | Cytophagales | Cyclobacteriaceae | Algoriphagus | 3 | 9 | 68.6 | 31.4 |
| 3eb9845cd243bc2cfcdbed443da59d4f | Prokaryote | Bacteroidota | Bacteroidia | Bacteroidales | VadinHA17 | SR-FBR-E99 | 2 | 9 | 0.0 | 100.0 |
| 0961323a97c36376c64a18274d521411 | Eukaryota | Ochrophyta | Bacillariophyta | Bacillariophyta_X | Araphid-pennate | NA_Araphid-pennate | 2 | 7 | 41.3 | 58.7 |
| ed9e188eff960b30e1ac074ca08370bb | Prokaryote | Proteobacteria | Alphaproteobacteria | Sphingomonadales | Sphingomonadaceae | Sphingomonas | 1 | 16 | 81.8 | 18.2 |
| 1acd1194f10b63af68420330712ea31f | Prokaryote | Proteobacteria | Alphaproteobacteria | Sphingomonadales | Sphingomonadaceae | Sphingomonas | 4 | 16 | 81.8 | 18.2 |
| e300ddf14f7cfd072b21591b4780924e | Eukaryota | Conosa | Variosea | Variosea_X | Schizoplasmodiids | Ceratiomyxella | 1 | 4 | 80.5 | 19.5 |
| Cluster 3 | | | | | | | | | | |
| 4b96b78bdad132c42872b483e056f04e | Eukaryota | Ochrophyta | Chrysophyceae | Chrysophyceae_X | Chrysophyceae_Cluster-D | Chrysophyceae_Cluster-D_X | 2 | 7 | 100.0 | 0.0 |
| 13cc45e417c97cc67ba8761f6a7f95c9 | Prokaryote | Cyanobacteria | Cyanobacteriia | Elainellales | Elainellaceae | NA_Elainellaceae | 1 | 11 | 89.6 | 10.4 |
| 99a00fa53f83a5613cf8df8f96056548 | Eukaryota | Ochrophyta | Bacillariophyta | Bacillariophyta_X | Raphid-pennate | NA_Raphid-pennate | 4 | 7 | 100.0 | 0.0 |
| cb36756419ecb275b1d0d4a1e8d06bdd | Eukaryota | Ochrophyta | Bacillariophyta | Bacillariophyta_X | Raphid-pennate | Nitzschia | 3 | 7 | 84.1 | 15.9 |
| 4205e76064917ac2c2c46f16652142c0 | Eukaryota | Ochrophyta | Bacillariophyta | Bacillariophyta_X | Raphid-pennate | Nitzschia | 4 | 7 | 100.0 | 0.0 |
| ccfe8681b9a8b0b0fd93443ae5299f71 | Prokaryote | Bacteroidota | Bacteroidia | Flavobacteriales | Flavobacteriaceae | Flavobacterium | 1 | 9 | 73.2 | 26.8 |
| 6e5ae7e2ddd7e7c5f97d5f239af7cd1a | Eukaryota | Ciliophora | Spirotrichea | Hypotrichia | Oxytrichidae | Oxytricha | 1 | 3 | | |
| 3a83ee6f1a20f77445081d315df96bff | Prokaryote | Cyanobacteria | Cyanobacteriia | Phormidesmiales | Phormidesmiaceae | Nodosilinea | 4 | 11 | 100.0 | 0.0 |
| 2a2fd1e0ae3b1ea9d6d7555e8f007594 | Prokaryote | Proteobacteria | Alphaproteobacteria | Sphingomonadales | Sphingomonadaceae | NA_Sphingomonadaceae | 1 | 16 | 100.0 | 0.0 |
| 0ec983f3026acebd13724448202a7036 | Prokaryote | Proteobacteria | Alphaproteobacteria | Rhodobacterales | Rhodobacteraceae | NA_Rhodobacteraceae | 1 | 16 | 100.0 | 0.0 |
| 39bb2b1b2f7a5f543f20877040c3a0f0 | Prokaryote | Delongbacteria | UBA4055 | UBA4055 | UBA4055 | UBA4055 | 1 | 12 | 68.8 | 31.2 |
| 9b90317f485a2c4ae2c7360673d0b6bf | Prokaryote | Bacteroidota | Bacteroidia | Chitinophagales | Chitinophagaceae | Phnomibacter | 3 | 9 | 100.0 | 0.0 |
| Cluster 4 | | | | | | | | | | |
| 5b1d2e16ff623ca86fa8eba17d767269 | Eukaryota | Ochrophyta | Chrysophyceae | Chrysophyceae_X | Chrysophyceae_Cluster-B2 | NA_Chrysophyceae_Cluster-B2 | 1 | 7 | 36.9 | 63.1 |
| 6337c7a4c82da992142769b354699be2 | Prokaryote | Proteobacteria | Alphaproteobacteria | Sphingomonadales | Sphingomonadaceae | Erythrobacter | 1 | 16 | 81.2 | 18.8 |
| 537eae19c81d46fe5c0643ecf65df809 | Eukaryota | Chlorophyta | Chlorophyceae | Chlamydomonadales | Chlamydomonadales_X | Chlamydomonas | 1 | 2 | 51.0 | 49.0 |
| 2fae73ba8f647d54aa8087c06681299b | Eukaryota | Ochrophyta | Bacillariophyta | Bacillariophyta_X | Raphid-pennate | Raphid-pennate_X | 3 | 7 | 100.0 | 0.0 |
| 3f784ab9fd75b6558b09b4f994af3c0b | Prokaryote | Proteobacteria | Alphaproteobacteria | Sphingomonadales | Sphingomonadaceae | Chakrabartia | 3 | 16 | 70.6 | 29.4 |
| bb2d2f5cced962f603ac2f00e8726bfa | Prokaryote | Gemmatimonadota | Gemmatimonadetes | Gemmatimonadales | Gemmatimonadaceae | Gemmatimonas | 2 | 15 | 100.0 | 0.0 |
| Cluster 5 | | | | | | | | | | |
| bcc1107831e3bccc7dadc6ad276481ea | Eukaryota | Ochrophyta | Bacillariophyta | Bacillariophyta_X | Raphid-pennate | NA_Raphid-pennate | 3 | 7 | 100.0 | 0.0 |
| bb1059351a482907ba7f6c8c10f24a5b | Prokaryote | Spirochaetota | Spirochaetia | Treponematales | Termitinemataceae | Treponema_G | 1 | 17 | 100.0 | 0.0 |
| ed306b4f6286d6042a3263e27f653d11 | Eukaryota | Ochrophyta | Bacillariophyta | Bacillariophyta_X | Raphid-pennate | Raphid-pennate_X | 5 | 7 | 100.0 | 0.0 |
| ba50bb8de30ea52954ab7184b4fd6f92 | Eukaryota | Ochrophyta | Bacillariophyta | Bacillariophyta_X | Raphid-pennate | Navicula | 1 | 7 | 100.0 | 0.0 |
| e7b3da6ef445a110ee3a03fc9f81ce7a | Prokaryote | Bacteroidota | Bacteroidia | Bacteroidales | VadinHA17 | NA_VadinHA17 | 2 | 9 | 100.0 | 0.0 |
| 198580222aca1797ea3b6aa396b170ab | Prokaryote | Proteobacteria | Gammaproteobacteria | Burkholderiales | Burkholderiaceae | NA_Burkholderiaceae | 1 | 16 | 78.1 | 21.9 |
| 5d121d7ec84a162e1c07a5c341695148 | Eukaryota | Cercozoa | Endomyxa | Vampyrellida | NA_Vampyrellida | NA_Vampyrellida | 1 | 1 | 85.6 | 14.4 |
| 45a51a823bf42e79bd642f159cbc3370 | Prokaryote | Proteobacteria | Alphaproteobacteria | Sphingomonadales | Sphingomonadaceae | Erythrobacter | 1 | 16 | 69.6 | 30.4 |
| f9ec4d3cac90588134b16a966e803a14 | Prokaryote | Bacteroidota | Bacteroidia | Cytophagales | Cyclobacteriaceae | JAAUQL01 | 2 | 9 | 67.7 | 32.3 |
| 034d421d0a95de4046e4eb0214e5e9a9 | Prokaryote | Cyanobacteria | Cyanobacteriia | Cyanobacteriales | NA_Cyanobacteriales | NA_Cyanobacteriales | 3 | 11 | 100.0 | 0.0 |
| Cluster 6 | | | | | | | | | | |
| 9bab90e5604ee2eaab627bcadae5cb6f | Eukaryota | Ochrophyta | Bacillariophyta | Bacillariophyta_X | Araphid-pennate | NA_Araphid-pennate | 1 | 7 | 51.4 | 48.6 |
| 633db22e72f80415804de79aa3389624 | Prokaryote | Cyanobacteria | Cyanobacteriia | Pseudanabaenales | seudanabaenaceae | Pseudanabaena | 1 | 11 | 100.0 | 0.0 |
| c094c0c315fcc34f4f6d3bb71590d864 | Eukaryota | Cercozoa | Endomyxa | Vampyrellida | NA_Vampyrellida | NA_Vampyrellida | 2 | 1 | 100.0 | 0.0 |
| a42acbf774d54c48c3a415db9f0df57f | Prokaryote | Proteobacteria | Gammaproteobacteria | Burkholderiales | Burkholderiaceae | Paucibacter_A | 3 | 16 | 100.0 | 0.0 |
| b854c09d7116c9f0bfc5a65a973045c9 | Eukaryota | Ochrophyta | Bacillariophyta | Bacillariophyta_X | Araphid-pennate | NA_Araphid-pennate | 1 | 7 | 100.0 | 0.0 |
| Cluster 7 | | | | | | | | | | |
| 0ac506096f3f5c210c5c38e4f8bf368d | Eukaryota | Ochrophyta | Bacillariophyta | Bacillariophyta_X | Raphid-pennate | Rhopalodia | 1 | 7 | 100.0 | 0.0 |
| 92a88ba031df04cfd8211df246df4949 | Eukaryota | Ochrophyta | Bacillariophyta | Bacillariophyta_X | Raphid-pennate | Rhopalodia | 2 | 7 | 100.0 | 0.0 |
| c458cc5b5c5c876b94df04ce9bbd1d40 | Prokaryote | Bacteroidota | Bacteroidia | Chitinophagales | Saprospiraceae | NA_Saprospiraceae | 1 | 9 | 100.0 | 0.0 |
| db16ac2b93eb226a1ee4d137de1fa775 | Prokaryote | Proteobacteria | Gammaproteobacteria | Pseudomonadales | Halieaceae | Chromatocurvus | 2 | 16 | 100.0 | 0.0 |
| Cluster 8 | | | | | | | | | | |
| 8f8e9b33207e42d5ece133221c5c0243 | Eukaryota | Ochrophyta | Bacillariophyta | Bacillariophyta_X | Raphid-pennate | NA | 1 | 7 | 0.0 | 100.0 |
| c2256795315c4da2c33510434c4fb78e | Prokaryote | Cyanobacteria | Cyanobacteriia | Cyanobacteriales | NA | NA | 1 | 11 | 8.8 | 91.3 |
| a8a2097de09e2fb9bc806a8a0136cfec | Eukaryota | Ochrophyta | Bacillariophyta | Bacillariophyta_X | Raphid-pennate | Raphid-pennate_X | 2 | 7 | 11.8 | 88.2 |
| b1038a3212d3975a5a55e0a7331e7112 | Prokaryote | Proteobacteria | Gammaproteobacteria | Burkholderiales | SG8-39 | SCGC-AG-212-J23 | 3 | 16 | 0.0 | 100.0 |
| aa75061450f48bf5278a56a18d87536c | Eukaryota | Ochrophyta | Bacillariophyta | Bacillariophyta_X | Raphid-pennate | Raphid-pennate_X | 1 | 7 | 0.0 | 100.0 |
| Cluster 9 | | | | | | | | | | |
| 9bb07d58b4a3da084df1a538ef75b91c | Eukaryota | Ochrophyta | Bacillariophyta | Bacillariophyta_X | Raphid-pennate | Pseudo-nitzschia | 1 | 7 | 52.4 | 47.6 |
| 99d54032e7b70bb0ccb5897ff95f78f8 | Prokaryote | Chloroflexota | Anaerolineae | Anaerolineales | EnvOPS12 | NA_EnvOPS12 | 1 | 10 | 34.0 | 66.0 |
| edff657783390e2148c4c5290f60cb87 | Eukaryota | Ochrophyta | Bacillariophyta | Bacillariophyta_X | Raphid-pennate | NA_Raphid-pennate | 2 | 7 | 41.9 | 58.1 |
| 623109c69c8e19c5c858e8f37f21b282 | Prokaryote | Proteobacteria | Gammaproteobacteria | Halothiobacillales | Halothiobacillaceae | Guyparkeria | 2 | 16 | 45.1 | 54.9 |
| Cluster 10 | | | | | | | | | | |
| 68f6d8297618f721959e1b947ff9f493 | Prokaryote | Bacteroidota | Ignavibacteria | Ignavibacteriales | Ignavibacteriaceae | IGN2 | 1 | 9 | 21.5 | 78.5 |
| 29cb722754737b032bafc5794ebd109c | Eukaryota | Ochrophyta | Bacillariophyta | Bacillariophyta_X | Araphid-pennate | NA_Araphid-pennate | 2 | 7 | 34.0 | 66.0 |
| c05bfbfbebd2f20c9bb0c15c43ebf470 | Eukaryota | Ochrophyta | Bacillariophyta | Bacillariophyta_X | Araphid-pennate | NA_Araphid-pennate | 1 | 7 | 33.3 | 66.7 |
| c015ffcfa7cf8c1d0703d29df0e586ca | Prokaryote | Proteobacteria | Gammaproteobacteria | Chromatiales | Chromatiaceae | Thiocapsa | 2 | 16 | 40.6 | 59.4 |
| Cluster 11 | | | | | | | | | | |
| 59a199c17f7338f3cc61f94c3a5edced | Eukaryota | Fungi | Cryptomycota | Cryptomycotina | Cryptomycotina_X | Cryptomycotina_XX | 2 | 5 | 62.6 | 37.4 |
| eab1706e57a19a653991bd7bc4973f53 | Prokaryote | Proteobacteria | Alphaproteobacteria | Rhodobacterales | Rhodobacteraceae | EhC02 | 1 | 16 | 62.5 | 37.5 |
| f79e0a19dd104445744f27b95585639b | Prokaryote | Bacteroidota | Bacteroidia | Bacteroidales | Prolixibacteraceae | Draconibacterium | 1 | 9 | 0.0 | 100.0 |
| 4fbbdf1733d05a2fbad89a5fd7a924ce | Prokaryote | Proteobacteria | Gammaproteobacteria | Burkholderiales | Burkholderiaceae | NA_Burkholderiaceae | 1 | 16 | 71.2 | 28.8 |
| 6362dd6ffaf6c487705e4b4c2ddebb81 | Prokaryote | Cyanobacteria | Cyanobacteriia | Cyanobacteriales | Nostocaceae | Dolichospermum | 2 | 11 | 74.5 | 25.5 |
| 4926b707b8249d90aa3aeea012bfd88a | Eukaryota | Ochrophyta | Bacillariophyta | Bacillariophyta_X | Raphid-pennate | NA_Raphid-pennate | 3 | 7 | 42.2 | 57.8 |
| Cluster 12 | | | | | | | | | | |
| 8745767fb4dcc3282e40c3528e9aa00b | Prokaryote | Bacteroidota | Bacteroidia | Cytophagales | Cyclobacteriaceae | Algoriphagus | 1 | 9 | 68.2 | 31.8 |
| 240efcb93a1b2639b0ad9687b5ddf211 | Eukaryota | Ochrophyta | Bacillariophyta | Bacillariophyta_X | Raphid-pennate | Raphid-pennate_X | 3 | 7 | 53.4 | 46.6 |
| b00686e4110c3fe130eb874df763ddd3 | Prokaryote | Gemmatimonadota | Gemmatimonadetes | Gemmatimonadales | Gemmatimonadaceae | JAABRT01 | 1 | 15 | 65.7 | 34.3 |
| c13a9ef671189f49d0cc0b7d96478097 | Prokaryote | Gemmatimonadota | Gemmatimonadetes | Gemmatimonadales | Gemmatimonadaceae | Gemmatimonas | 1 | 15 | 63.0 | 37.0 |
| Cluster 13 | | | | | | | | | | |
| d2ceb513e4e34e7c14d0e5a88aa7ca99 | Prokaryote | Proteobacteria | Alphaproteobacteria | Caulobacterales | TH1-2 | Aquidulcibacter | 1 | 16 | 100.0 | 0.0 |
| 25689b3975a59e0af4337732e06869f2 | Eukaryota | Ciliophora | Nassophorea | Nassophorea_X | Nassulida | NA_Nassulida | 3 | 3 | 0.0 | 100.0 |
| 20bf68f2871a729e181e20b1a32bffa9 | Prokaryote | Bacteroidota | Bacteroidia | Bacteroidales | UBA7960 | SKVR01 | 2 | 9 | 18.1 | 81.9 |
| 14966980255fe5c334bc3564209cf4de | Prokaryote | Bacteroidota | Ignavibacteria | Ignavibacteriales | Ignavibacteriaceae | IGN3 | 1 | 9 | 29.0 | 71.0 |
| 1bb84f2078088527efbf501cd555230a | Prokaryote | Proteobacteria | Alphaproteobacteria | Sphingomonadales | Sphingomonadaceae | Erythrobacter | 1 | 16 | 100.0 | 0.0 |
| 20e1edd2ef7180c06bf17d907584ca38 | Prokaryote | Desulfobacterota | Desulfobacteria | Desulfobacterales | Desulfatirhabdiaceae | Desulfatirhabdium | 1 | 13 | 32.6 | 67.4 |
| fdc88d68afc8bbd69ba1d3b6ff6d1466 | Eukaryota | Ciliophora | Spirotrichea | Hypotrichia | Oxytrichidae | Oxytricha | 3 | 3 | 84.1 | 15.9 |
| Cluster 14 | | | | | | | | | | |
| 701383e3282f2e0868b5b8bb985e0c01 | Eukaryota | Ochrophyta | Bacillariophyta | Bacillariophyta_X | Raphid-pennate | Raphid-pennate_X | 3 | 7 | 71.2 | 28.8 |
| 2f588d3417afc605ce8162cc72694f09 | Prokaryote | Proteobacteria | Gammaproteobacteria | Burkholderiales | Burkholderiaceae | NA_Burkholderiaceae | 1 | 16 | 71.5 | 28.5 |
| b1be65d2068e4dbd47a0394db0e57960 | Prokaryote | Proteobacteria | Alphaproteobacteria | Sphingomonadales | Sphingomonadaceae | NA_Sphingomonadaceae | 1 | 16 | 83.7 | 16.3 |
| 177e01ce2fc91e5731208079ff019355 | Prokaryote | Gemmatimonadota | Gemmatimonadetes | Gemmatimonadales | GWC2-71-9 | JACDDX01 | 1 | 15 | 79.7 | 20.3 |
| Cluster 15 | | | | | | | | | | |
| bce73e8a2f692c7d435d5e81d3d070b6 | Eukaryota | Ochrophyta | Chrysophyceae | Chrysophyceae_X | Chrysophyceae_Cluster-D | Chrysophyceae_Cluster-D_X | 1 | 7 | 0.0 | 100.0 |
| e3f806f6ebc7442f7745e1aedcadedb6 | Eukaryota | Fungi | Cryptomycota | Cryptomycotina | Cryptomycotina_X | Cryptomycotina_XX | 1 | 5 | 45.8 | 54.2 |
| 597107857d7e4ade0da855f7f4ac8961 | Eukaryota | Ochrophyta | Bacillariophyta | Bacillariophyta_X | Raphid-pennate | NA_Raphid-pennate | 3 | 7 | 45.2 | 54.8 |
| 0ab741bd7068f236fb1f6100bc48387b | Prokaryote | Bacteroidota | Bacteroidia | Flavobacteriales | Flavobacteriaceae | Flavobacterium | 1 | 9 | 59.2 | 40.8 |
| 93286f405ddbd0dcc60049eaadfd51f1 | Prokaryote | Firmicutes | Bacilli | Erysipelotrichales | Erysipelotrichaceae | UBA6182 | 3 | 14 | 26.3 | 73.7 |
| ee5672026ab8c5137a082457f814b10b | Prokaryote | Bacteroidota | Bacteroidia | Cytophagales | Cyclobacteriaceae | NA_Cyclobacteriaceae | 1 | 9 | 67.7 | 32.3 |
| Cluster 16 | | | | | | | | | | |
| d0d1f1163ecd8c59d10ef4d6f413271e | Prokaryote | Firmicutes_F | Halanaerobiia | Halanaerobiales | Halanaerobiaceae | Halanaerobium | 1 | 14 | 66.2 | 33.8 |
| 1da19ff2992959045782d88a9cb5f9ce | Eukaryota | Ochrophyta | Bacillariophyta | Bacillariophyta_X | Raphid-pennate | Navicula | 3 | 7 | 86.6 | 13.4 |
| 8db5591bf11fe1b03e5b91439426eca2 | Prokaryote | Proteobacteria | Alphaproteobacteria | Rhodobacterales | Rhodobacteraceae | NA_Rhodobacteraceae | 1 | 16 | 79.3 | 20.7 |
| d7824c14b705570877fb2c8a15f4784b | Eukaryota | Ochrophyta | Bacillariophyta | Bacillariophyta_X | Raphid-pennate | Raphid-pennate_X | 1 | 7 | 40.0 | 60.0 |
| 20b47382ea6a116db68454aa9cc68b2f | Prokaryote | Desulfobacterota | Desulfobacteria | Desulfobacterales | Desulfobacteraceae | Desulfotignum | 3 | 13 | 60.6 | 39.4 |
| 4303d0563dc9b60c95ebb6667ea85e23 | Prokaryote | Desulfobacterota | Desulfobacteria | Desulfobacterales | Desulfobacteraceae | Desulfotignum | 2 | 13 | 47.9 | 52.1 |
| 6623ec873cc7c30bff146583539628b7 | Eukaryota | Cercozoa | Endomyxa | Vampyrellida | NA_Vampyrellida | NA_Vampyrellida | 3 | 1 | 58.2 | 41.8 |
| ecee79c5f0c1973beffc33ba2dcc8cba | Prokaryote | Proteobacteria | Gammaproteobacteria | Ectothiorhodospirales | Thioalkalivibrionaceae | Thioalkalivibrio_B | 1 | 16 | 78.8 | 21.2 |
| 8499ea48ca5d14bb0809c5254d8b6664 | Eukaryota | Ochrophyta | Bacillariophyta | Bacillariophyta_X | Raphid-pennate | Raphid-pennate_X | 2 | 7 | 52.9 | 47.1 |
| 0c4bdb225385e3aa7450742766b59d12 | Prokaryote | Desulfobacterota | Desulfobacteria | Desulfobacterales | Desulfobacteraceae | Desulfotignum | 1 | 13 | 57.9 | 42.1 |
| Cluster 17 | | | | | | | | | | |
| 2b8ea7b003234e87949f4961abfe14c2 | Eukaryota | Ochrophyta | Bacillariophyta | Bacillariophyta_X | Araphid-pennate | NA_Araphid-pennate | 2 | 7 | 46.0 | 54.0 |
| 363b85e6107a7b76e7746caad3c6245e | Prokaryote | Bacteroidota | Bacteroidia | Bacteroidales | VadinHA17 | SR-FBR-E99 | 1 | 9 | 26.6 | 73.4 |
| 2c87c68a764a0676138117fe367892ad | Eukaryota | Ochrophyta | Chrysophyceae | Chrysophyceae_X | Chrysophyceae_Cluster-C | Chrysophyceae_Cluster-C_X | 1 | 7 | 57.4 | 42.6 |
| cdcf16b7c5bd37921f3ca53136e8a8ee | Prokaryote | Gemmatimonadota | Gemmatimonadetes | Gemmatimonadales | Gemmatimonadaceae | Gemmatimonas | 2 | 15 | 86.4 | 13.6 |
| Cluster 18 | | | | | | | | | | |
| d74e0521f607b92af1021f1a1d12074d | Prokaryote | Desulfobacterota_I | Desulfovibrionia | Desulfovibrionales | Desulfomicrobiaceae | Desulfomicrobium | 1 | 13 | 70.9 | 29.1 |
| 1e815f1b77723124c9314d1e6a312388 | Prokaryote | Bacteroidota | Bacteroidia | Bacteroidales | JAAYUG01 | JAAYUG01 | 1 | 9 | 100.0 | 0.0 |
| 91cf9e0333f1407780369ffa22ee0004 | Eukaryota | Sagenista | Labyrinthulomycetes | Labyrinthulida | Labyrinthulaceae | Labyrinthulaceae_ANT10_3 | 1 | 8 | 61.7 | 38.3 |
| 07e2404d756f6aa57ef7cb850adaeb8b | Eukaryota | Lobosa | Tubulinea | Nolandida | Nolandellidae | NA_Nolandellidae | 2 | 6 | 100.0 | 0.0 |
| 26594b2bbdf2d9f5c2e0428dac873bd8 | Eukaryota | Conosa | Variosea | Variosea_X | Schizoplasmodiids | Ceratiomyxella | 3 | 4 | 100.0 | 0.0 |
| 5e72d1f79a9ec7f4561fa09954fb9c2e | Eukaryota | Ciliophora | Oligohymenophorea | Scuticociliatia_2 | Scuticociliatia_2_X | Cyclidium_3 | 4 | 3 | 100.0 | 0.0 |
| e00dc8ce50f6b10adbf060a06c02696d | Prokaryote | Cyanobacteria | Cyanobacteriia | Elainellales | Elainellaceae | NA_Elainellaceae | 2 | 11 | 100.0 | 0.0 |
| e221d93660429da9ccbef0c2f1eaf700 | Prokaryote | Cyanobacteria | Cyanobacteriia | Elainellales | Elainellaceae | RECH01 | 1 | 11 | 100.0 | 0.0 |
| 037dc1c9a772870b6c6b78f84208f235 | Prokaryote | Proteobacteria | Alphaproteobacteria | Acetobacterales | Acetobacteraceae | Roseococcus | 1 | 16 | 100.0 | 0.0 |
| 3e1536ac3d29a27281ed523d9fa239ca | Prokaryote | Bacteroidota | Rhodothermia | Balneolales | Balneolaceae | NA_Balneolaceae | 1 | 9 | 100.0 | 0.0 |
| 2ff4a743b59b99e2886e87f77477791c | Prokaryote | Proteobacteria | Alphaproteobacteria | Rhodobacterales | Rhodobacteraceae | Yoonia | 3 | 16 | 77.9 | 22.1 |
| Cluster 19 | | | | | | | | | | |
| d797085c7c7eb45c99b6cb98a55e4d41 | Prokaryote | Bacteroidota | Bacteroidia | Cytophagales | Cyclobacteriaceae | ELB16-189 | 1 | 9 | 69.3 | 30.7 |
| d375de2d531e15ad51d52949a4161b99 | Eukaryota | Ochrophyta | Bacillariophyta | Bacillariophyta_X | Araphid-pennate | NA_Araphid-pennate | 2 | 7 | 67.3 | 32.7 |
| b14adabb49c72a6cc77bb814c7b87ec5 | Eukaryota | Ochrophyta | Bacillariophyta | Bacillariophyta_X | Raphid-pennate | NA_Raphid-pennate | 2 | 7 | 45.9 | 54.1 |
| d6283176213c91ffbabd14c929fb1ed2 | Prokaryote | Desulfobacterota | Desulfobacteria | Desulfobacterales | NA_Desulfobacterales | NA_Desulfobacterales | 2 | 13 | 30.0 | 70.0 |

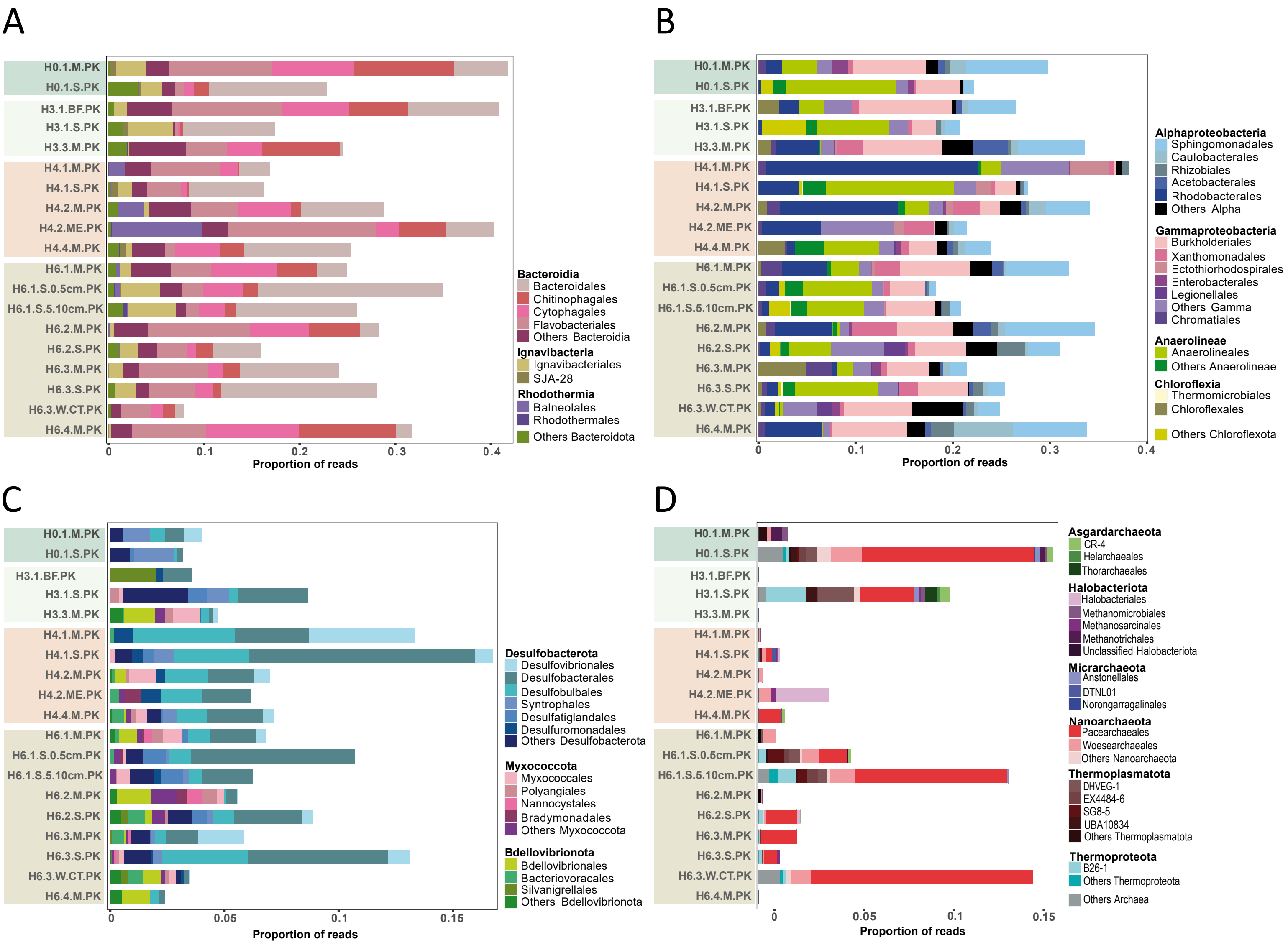


**Fig. S1**. Order-level taxonomic composition of selected high-rank prokaryotic taxa. A, Bacteroidetes. B, Proteobacteria (Alpha- and Gammaproteobacteria). C, Deltaproteobacteria. D, archaea.

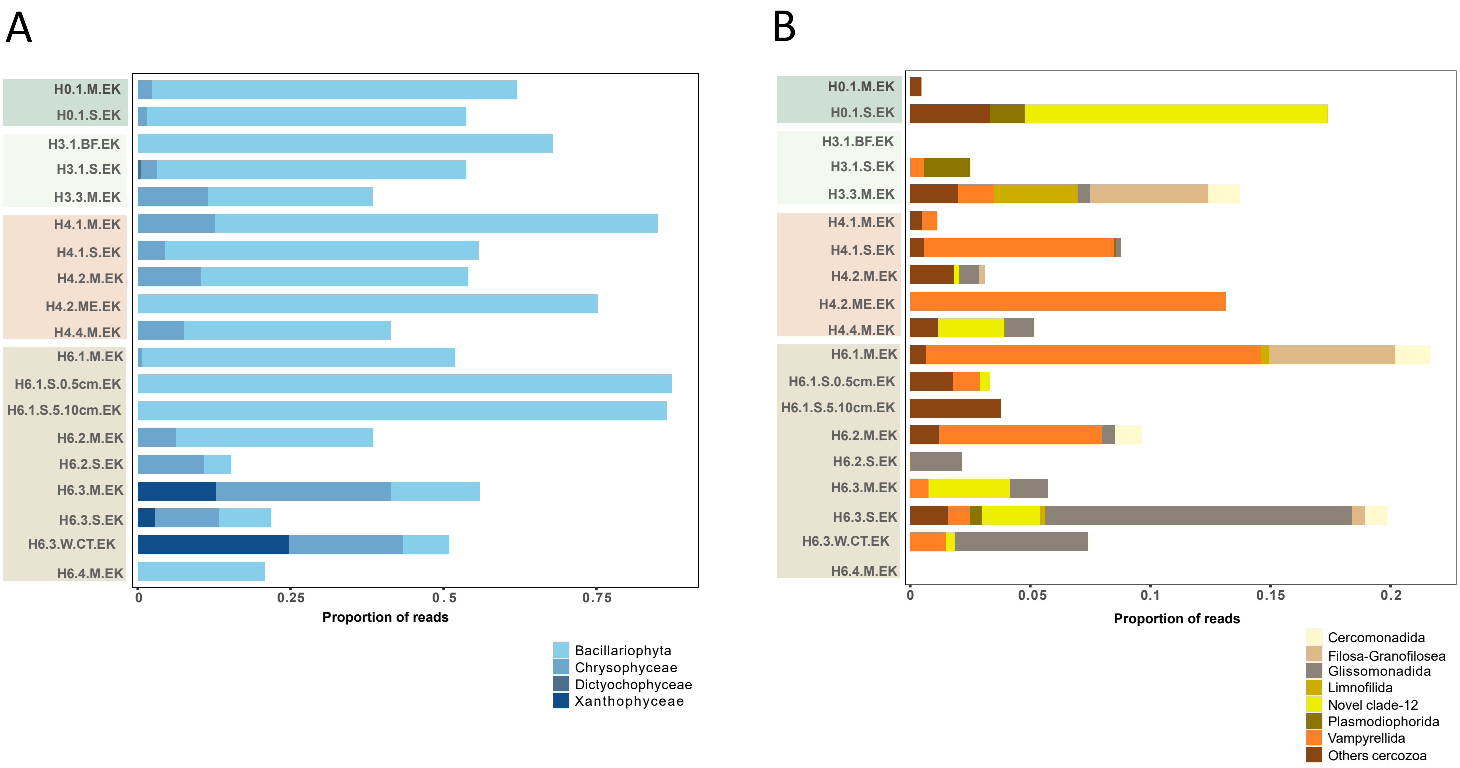


**Fig. S2**. Order-level taxonomic composition of selected high-rank eukaryotic taxa. A, Ochrophyta. B, Cercozoa

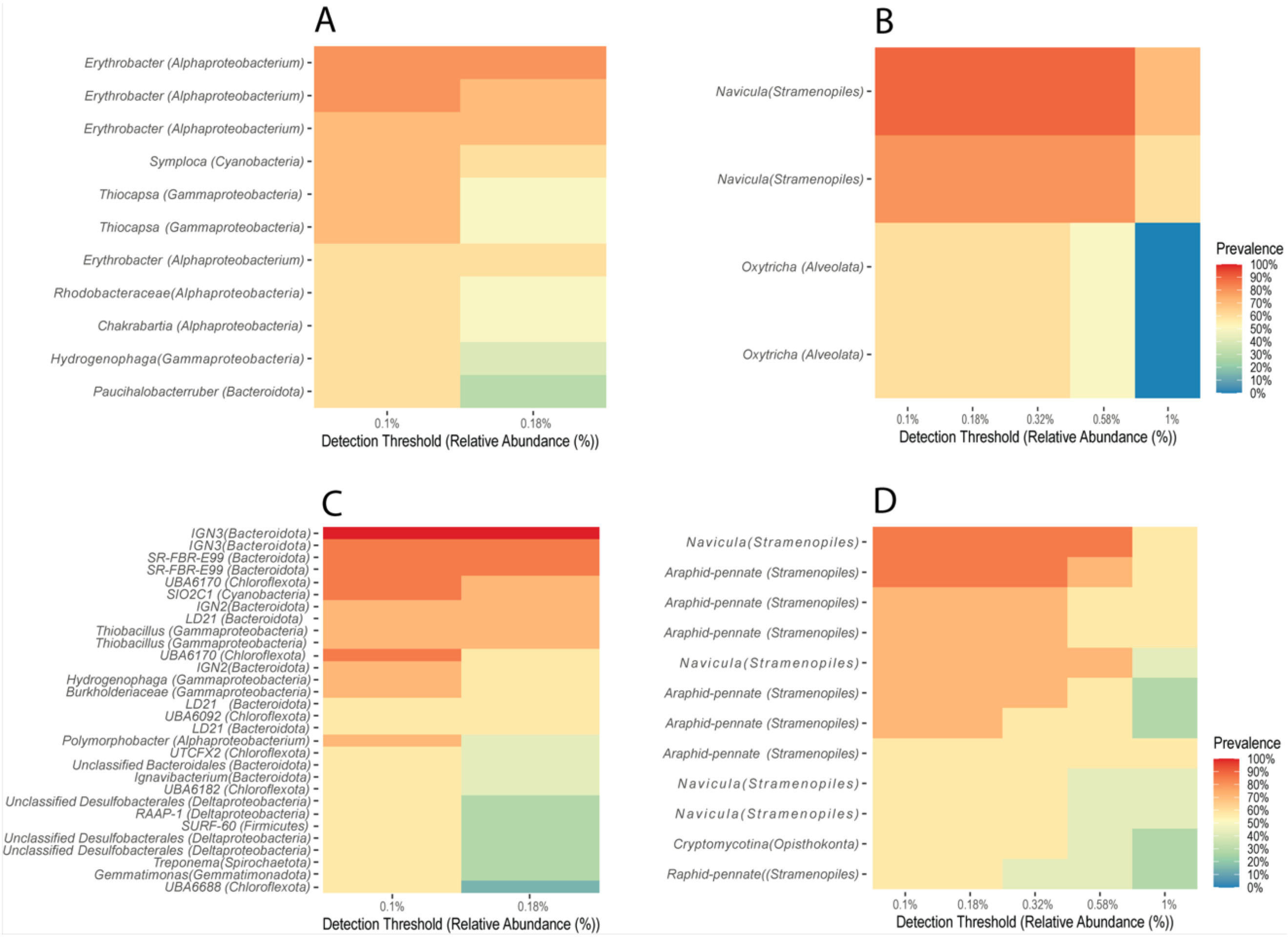


**Figure S3.** Composition of the prokaryotic and eukaryotic microbiomes core for microbial mats and sediment. A, prokaryotic core for microbial mats. B, eukaryotic cores for microbial mats. C, prokaryotic core for sediment samples. D, eukaryotic core for sediment samples.

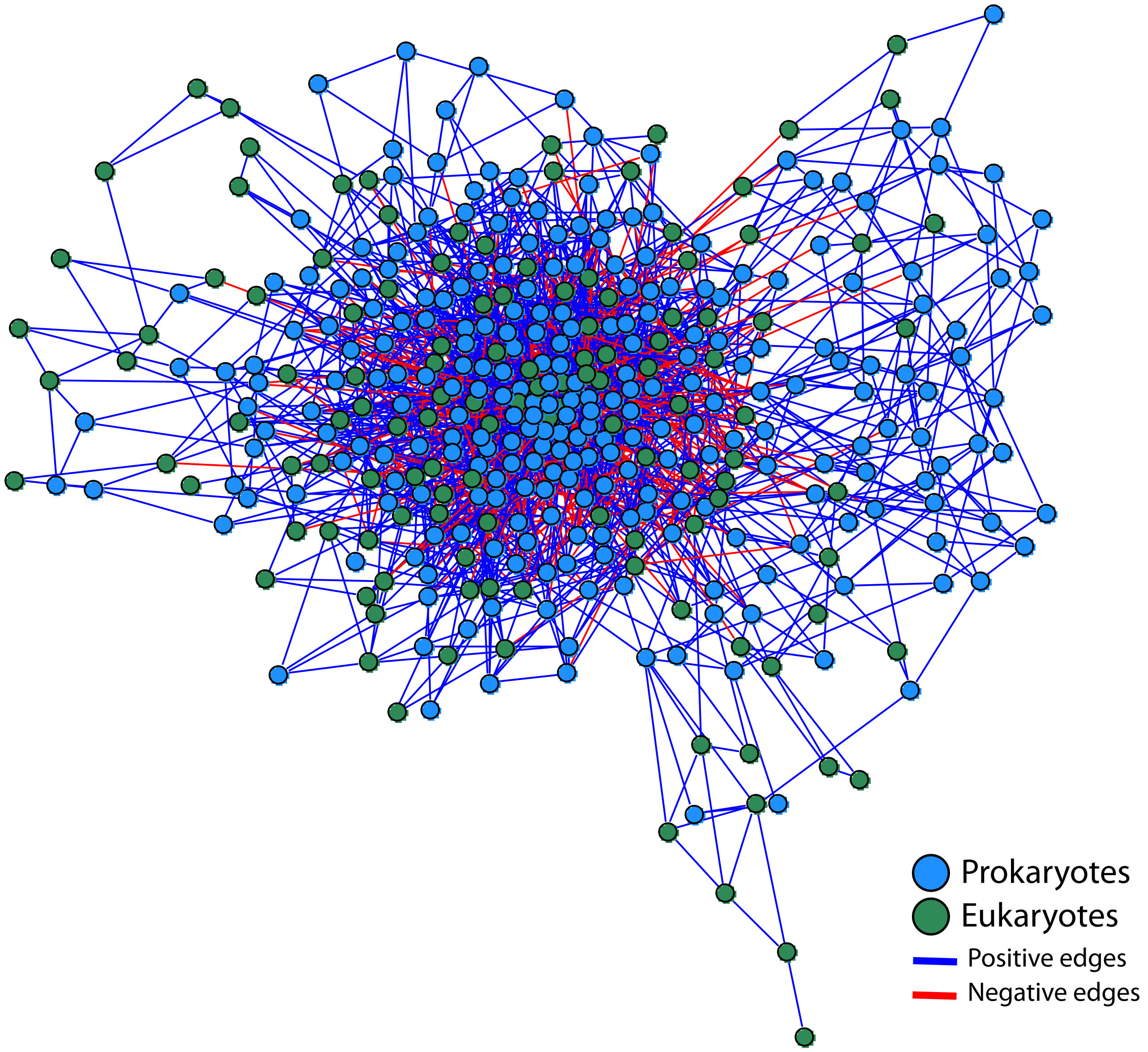


**Fig. S4**. Co-occurrence network of prokaryotic and eukaryotic members of mat and sediment samples from Salar de Huasco.
